# A membrane-free spot-plating protocol for *Agrobacterium*-mediated transformation of diverse yeasts

**DOI:** 10.64898/2026.03.02.708628

**Authors:** Matthew R. Incha, Matthew J. Szarzanowicz, Gina M. Geiselman, Amy Lanctot, Paul A. Adamczyk, Valentina E. Garcia, Mitchell G. Thompson, Patrick M. Shih, John M. Gladden, Di Liu

**Author notes:** Correspondence should be addressed to Di Liu.

## Abstract

*Agrobacterium*-mediated transformation (AMT) is a critical method for genetic manipulation of non-model fungi, yet it remains a laborious and inefficient technique. When cocultured in acetosyringone-supplemented induction medium, *Agrobacterium* transfers DNA directly into yeast cells using its virulence machinery. Membrane filters are commonly used to support the co-culture of yeast and *Agrobacterium* on agar plates, however some reports demonstrate that these filters are unnecessary for specific yeast species. Here we confirm across diverse budding yeasts that membrane filters are not necessary for effective AMT. Concentrating the cells via centrifugation and “spotting” the cell pellet directly onto the induction medium proved effective. This reduces hands-on time to 15 minutes and eliminates filter cost. In the oleaginous yeast, *Rhodotorula* toruloides, this simplified method increases transformation efficiency by 66% to 2,500 transformants per 10^6^ recipient cells. We further optimized the *Agrobacterium*:*Rhodotorula* cell ratio and culture resuspension volume to achieve more than 200,000 CFU per transformation representing a 2-3 fold improvement over previously implemented protocols. This spot-plating method was successfully applied to seven yeast species, including one for which genetic transformation has not previously been reported, *Botryozyma nematodophila*. This approach highlights the broad applicability of the spot-plating method across diverse yeast systems. Furthermore, this method could facilitate high-throughput transformation workflows that are critical for genome-scale functional studies.

## Introduction

Fungal transformation is critical for basic biological research and applied technical applications in pharmaceuticals, bioproduction, and food fermentation. The investigation and biotechnological use of these important organisms is often hindered by our inability to efficiently transform heterologous genetic material into these species. Unlike *Saccharomyces* spp. and *Komagataela* (*Pichia*) spp., many fungal species are recalcitrant to transformation via traditional methods of electroporation or chemical transformation, often requiring laborious protoplasting protocols (1). This makes *Agrobacterium-*mediated transformation (AMT) an advantageous method for introducing heterologous DNA to these species and making genetic knockouts.

AMT leverages the naturally pathogenic relationship between *A. tumefaciens* and plants. In a mechanism analogous to bacterial conjugation, *Agrobacterium* transmits DNA encoding oncogenes into plant cells inducing tumor formation, the characteristic symptom of crown-gall disease. The oncogenes and the virulence genes (*vir* genes) that enable their transmission are encoded on the tumor-inducing (Ti) plasmid of *Agrobacterium* (2,3). The oncogenes are located in a region called the transfer-DNA (T-DNA) and are flanked by defined DNA sequences known as the right border (RB) and left border (LB). Facilitated by the virulence proteins, the T-DNA sequence between these borders is excised, transferred into recipient cells, escorted to the nucleus, and integrated into the host genome (4).

Capitalizing on *Agrobacterium*’s plant-genome transformation capacity, plant biologists domesticated *Agrobacterium* by removing oncogenes from the Ti plasmids and porting the LB and RB sequences into smaller synthetic plasmids known as binary vectors (5– 7). By cloning transgenes between these conserved border sequences, biologists are able to stably introduce novel transgenic sequences directly into the plant genome, revolutionizing plant genetics and engineering (8). Later, AMT was implemented to transgenically modify fungal species (9). In this pivotal work of Bundock et al., *Saccharomyces cerevisiae* and *Agrobacterium* were mixed and added directly onto filters placed on an induction medium plate (9), a protocol presumably modified from standard practice for bacterial conjugation (10). However, since this first description, many groups have employed vacuum filtration of the coculture to embed larger quantities of cells onto the membrane (11).

While filter membranes are helpful for handling filamentous fungi after co-cultivation (12), the use of filters for budding-yeasts appears, at first glance, unnecessary. The additional step of vacuum filtration adds time, labor, batch-variability, and risk of contamination. Mitigating these challenges is important for the widespread adoption of this transformation method. Modifications to the protocol have been made to improve transformation efficiency, such as changing the choice of filter polymer and the method of embedding cells onto the filter (13). Despite this, only a few reported methods in *Cryptococcus* and *Rhodotorula* genetics studies bypass the filter altogether and spot the mixture of *Agrobacterium* and yeast directly onto the induction medium (14,15). Here, we demonstrate that a spot-plating method omitting filter membranes and vacuum filtration can be optimized and broadly deployed across diverse yeast species.

## Results and Discussion

### Spot plating improves transformation efficiency, reduces protocol time, and does not impact transformation outcomes compared to filtered transformations

In this study, we primarily focused on the yeast *Rhodotorula toruloides* IFO0880 (NBRC 0880/NRRL 1588), as we routinely achieve more than 50,000 colonies per transformation in this species using AMT with vacuum filtration. All transformations in this host were conducted with the binary vector plasmid, pVL_sR2rn_f14rg, containing the *R. graminis* WP1 TUB2 promoter driving a nourseothricin resistance gene, *nrsR*, and the P14 promoter, driving a codon optimized sfGFP (rg) (16).

The standard protocol is briefly described here: *Agrobacterium tumefaciens* strain EHA105 carrying a desired binary vector is cultured in LB to an OD of ∼1. The culture is then centrifuged, washed, and incubated overnight in an equal volume of induction medium. The target yeast is separately cultured to an OD of ∼1 in YPD (4.5E7 cells for *Rhodotorula*). One milliliter of yeast is pelletted, the YPD is aspirated and the pellet resuspended with one milliliter of acetosyringone-induced *Agrobacterium*. The *Agrobacterium*-yeast mixture is then pulled onto a 0.45 µm mixed cellulose ester filter under vacuum, and the filter is placed onto an induction medium plate for 4-5 days. After growth on the induction medium, the co-culture is then removed from the filter, resuspended in YPD, and culture dilutions are plated onto selective medium plates with the addition of beta-lactams (cefotaxime, carbenicillin, or timentin) to prevent *Agrobacterium* growth.

The process of vacuum-pulling and drying the mixed culture of yeast and *Agrobacterium* onto a filter membrane is incredibly laborious and time-consuming. With standard bench-top vacuum pumps (0.1-0.4 bar), this step can vary from 10 minutes to an hour for each filter, hindering higher-throughput transformation workflows. Using a single filtration apparatus, making a pooled library of 2 × 10^6^ transformants would require dozens of filters and hours of hands-on labor (17). To demonstrate this labor requirement, three researchers conducted and timed AMT of *Rhodotorula* using two distinct protocols using the same batches of *Agrobacterium* and *Rhodotorula*: the vacuum filtration protocol and a modified protocol wherein the cell mixture was added directly onto the induction medium. Two of the three researchers completed 6 replicates of the protocol including vacuum filtration in approximately three hours. One researcher had a fault in their vacuum apparatus that could not be identified, and they completed 6 filtration replicates in 5 hours. This extended period resulted in extensive contamination, and their data were unviable.

In contrast, the modified protocol that omits membrane filters and vacuum filtration took approximately 15 minutes to complete 6 replicates for all three researchers. This method simply requires centrifuging the mixture of *Rhodotorula* and *Agrobacterium*, aspirating the supernatant, resuspending the cell pellet, and pipetting the entire mixture as a droplet directly onto the induction medium plate (“spot-plating”). These plates were then left to dry for 10 minutes in a biosafety cabinet. After growth on the induction medium for 4-5 days, this protocol yields a spot of mixed cells approximately 1 to 2 cm in diameter.

Notably, the spot-plating method resulted in significantly more transformants than the vacuum filtration method. The vacuum filtration method yielded an average of 67,233 ± 14,269 CFU per transformation (range: 50,400–95,200) and an efficiency of 1,488 transformants per 10^6^ of starting *Rhodotorula* cells. The spot method produced 111,600 ± 19,782 CFU (range: 64,000–143,040) and an efficiency of 2,480 transformants per 10^6^ of starting *Rhodotorula* cells (Figure 1A). This represents a 66% increase in transformation efficiency for the spot-plating method relative to the standard vacuum filtration protocol (*p* < 0.001, Welch’s two-tailed *t*-test). This improvement was consistent across the transformation efficiencies obtained by both researchers, indicating the method’s reproducibility.

**Figure 1:**
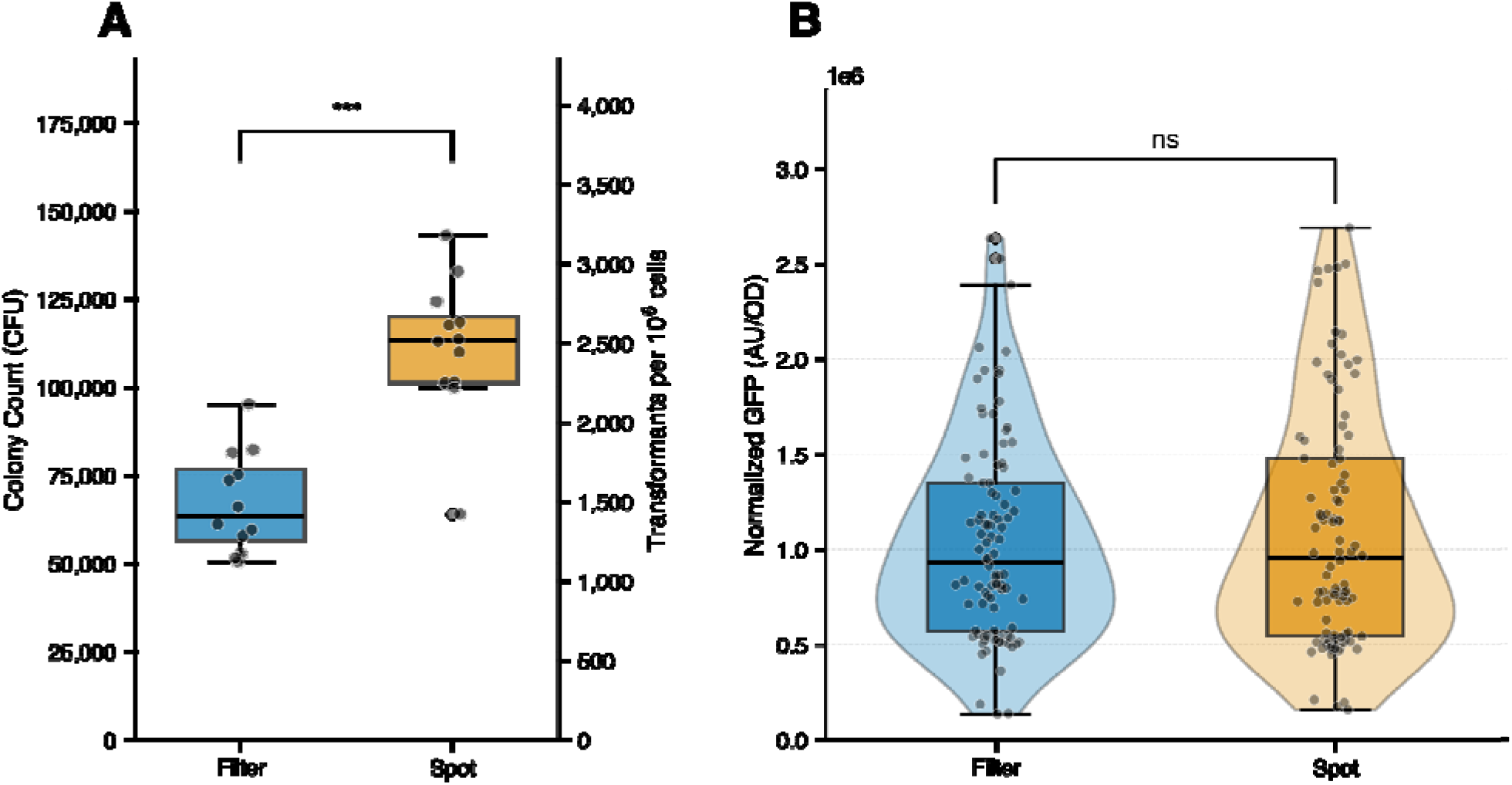
Comparison of filter and spot methods in *Rhodotorula toruloides*. **A.** Transformation results in terms of number of CFUs from the filter and spot plating methods (n = 12 filters and 12 spots, p = 0.000004 Welch’s t-test). **B**. Comparison of the transformation outcomes in terms of GFP intensity from randomly selected colonies from transformants in 1A (n = 96 per group, p = 0.5364 Welch’s t-test, p = 0.99 Mann-Whitney U). All transformations were conducted with the binary vector pVL_sR2rn_f14rg.

Transformation efficiency is not the only metric critical for a transformation protocol, as the genetic outcome of the transformation is also an essential criterion. To test if the transformation outcome was equivalent between the two protocols, we estimated the transgene copy number in 96 colonies from each method using fluorescence intensity as a proxy for GFP insertion events (18). Transformed colonies were cultured in YPD, and their fluorescence was measured using a plate-reader. The levels of fluorescence in these populations were indistinguishable, suggesting that both methods yielded similar copy numbers of transgenic integration events (Figure 1B).

### *Agrobacterium*:Yeast Ratio contributes to transformation efficiency

To further optimize the spot-plating method, we systematically tested how cell densities of *Rhodotorula* and *Agrobacterium* affect transformation efficiency. Cell quantities are reported as OD·mL (OD□□□ × volume in mL), where 1 OD·mL represents the cell content of 1 mL of culture at OD□□□ = 1. Cultures were retrieved at OD□□□ = 1 before use, and varying volumes were concentrated or diluted to achieve the indicated OD·mL values. Three cell densities (0.2, 1, and 5 OD·mL) for both species were tested across nine pairwise conditions. All 9 combinations produced viable transformants, but efficiency varied dramatically (34 to 1,594 CFU per 10^6^ *Rhodotorula* cells). The highest yield of transformants was achieved with the highest combined cell density (5 OD·mL each; mean: 218,333), representing a 3-fold improvement over the standard density (1 OD·mL each; mean: 71,750 CFU) (Figure 1A). Transformation efficiency was slightly lower at the highest cell density ratio compared to the standard density, though this difference was not statistically significant (Figure 2B). When the density of *Rhodotorula* cells was high (5 OD·mL), efficient transformation required proportionally higher densities of *Agrobacterium* (5 OD·mL) (Figure 2A,B). This may be attributed to reduced *Agrobacterium*-*Rhodotorula* cell-to-cell contacts due to occlusion by the high quantity of *Rhodotorula* cells.

**Figure 2:**
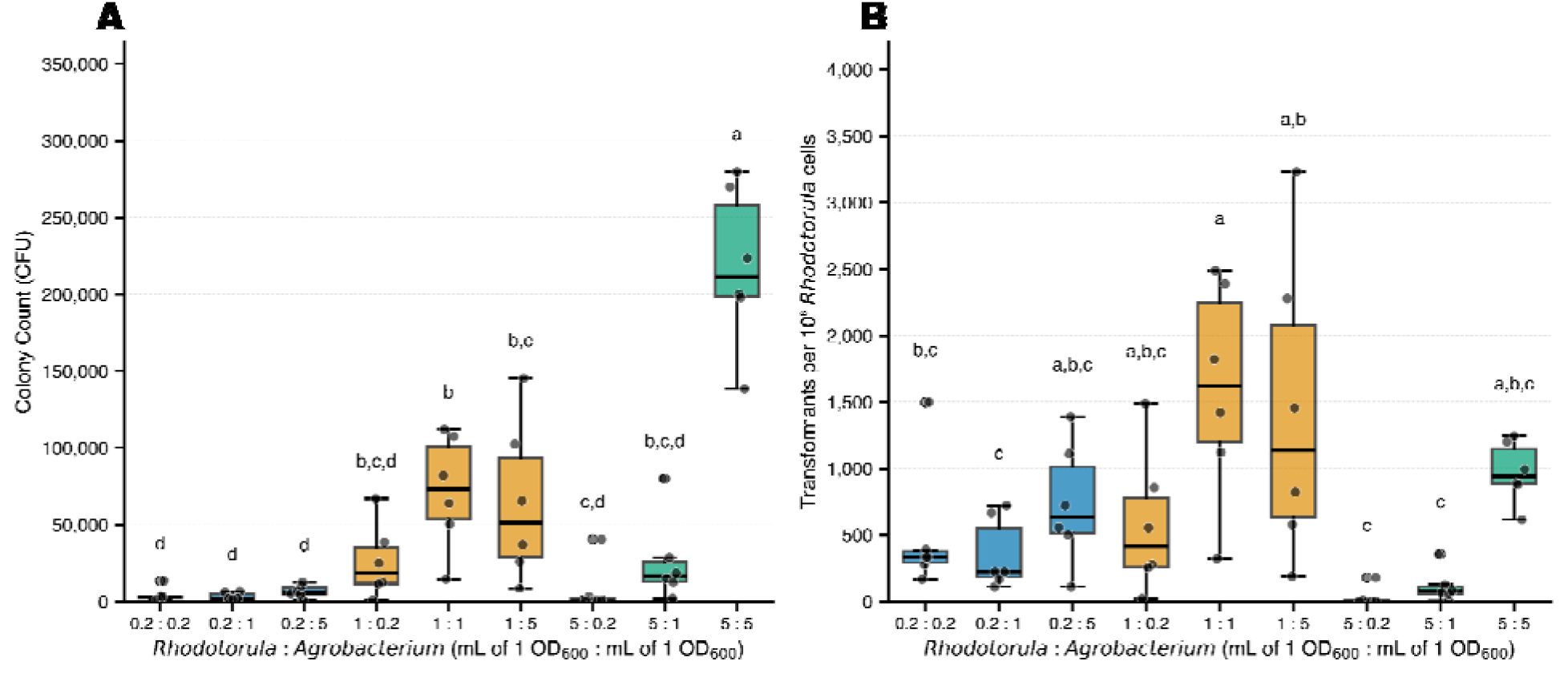
Cell ratio and spot volume effects on transformation efficiency. **A.** Total colonies per transformation with different proportions of *Agrobacterium* and *Rhodotorula* in terms of OD·mL. Letters above each group denote statistically distinct groups based on Tukey’s HSD post-hoc test (familywise error rate (FWER) = 0.05) following a significant one-way ANOVA (F = 30.56, p < 0.0001). Groups sharing a letter are not significantly different from one another. **B**. Transformation efficiency in terms of transformants per 10^6^ *Rhodotorula* cells with different proportions of *Agrobacterium* and *Rhodotorula*. Letters above each group denote statistically distinct groups based on Tukey’s HSD post-hoc test (FWER = 0.05) following a significant one-way ANOVA (F = 5.55, p < 0.0001). Groups sharing a letter are not significantly different from one another.

The size of the co-culture spot could additionally influence cell-to-cell contact. To evaluate this, we tested three resuspension volumes (20 μL, 75 μL, and 150 μL) using the standard OD·mL cell densities (1 OD·mL each). The 75 μL volume produced the highest colony counts (mean: 92,600 CFU), compared to 87,933 for 20 μL and 63,733 for 150 μL; however, these results were not statistically significant (Figure S1). The resulting spots are of increasing diameter as the volume increases. Upon drying, cell-cell contact remains sufficient for effective AMT across the range of volumes tested.

### Unfiltered spot-plating method is effective across species in diverse genera

To demonstrate broader applicability of the unfiltered spot-plating method for other yeast species, we first compared the filter and spot-plating methods using *Saccharomyces cerevisiae* and *Kluyveromyces marxianus. S. cerevisiae* was the first fungus transformed via AMT, while *K. lactis*, a close relative of *K. marxianus*, was also among the earliest (9,19). For *S. cerevisiae*, we constructed a plasmid (pVR_sATk_fS3sg_HR) containing the *Ashbya gossypii* TEF1 promoter driving a G418 resistance cassette, the *S. cerevisiae* TDH3 promoter driving sg12 GFP expression, and homology arms targeting the *tau* locus of chromosome XVI. Transformation of *K. marxianus* was conducted with pVR_sATk_fS3sg, an otherwise identical plasmid without homology arms. *K. marxianus* strains have demonstrated preference for NHEJ mechanisms similar to many environmental yeast strains making this non-targeting plasmid an effective test-case (20,21). Contrary to the results in *Rhodotorula*, the spot-plating method yielded fewer colonies for both hosts. Though this decrease was not statistically significant for *S. cerevisiae* (p > 0.05) but was highly significant for *K. marxianus* (p < 0.001) (Figure 3). *S. cerevisiae* yielded 99,400 ± 39,972 and 66,400 ± 19604 transformants in the filter and spot-plating methods, respectively. *K. marxianus* yielded 3,860 ± 811 and 880 ± 421 transformants in the filter and spot-plating methods respectively. However, the 5 replicates for each host required between 15 to 45 minutes for each filter to be pulled and dried (3 hours total), while the spot-plating was conducted in parallel and finished in 30 minutes (accounting for dry-time in a biosafety cabinet). The reduced efficiency when spot-plating these hosts could be due to any number of morphological differences relative to *Rhodotorula toruloides*. Notably, *Rhodotorula* cell walls contain distinct exopolysaccharides rich in mannose that may positively affect *Agrobacterium* adhesion (22).

**Figure 3:**
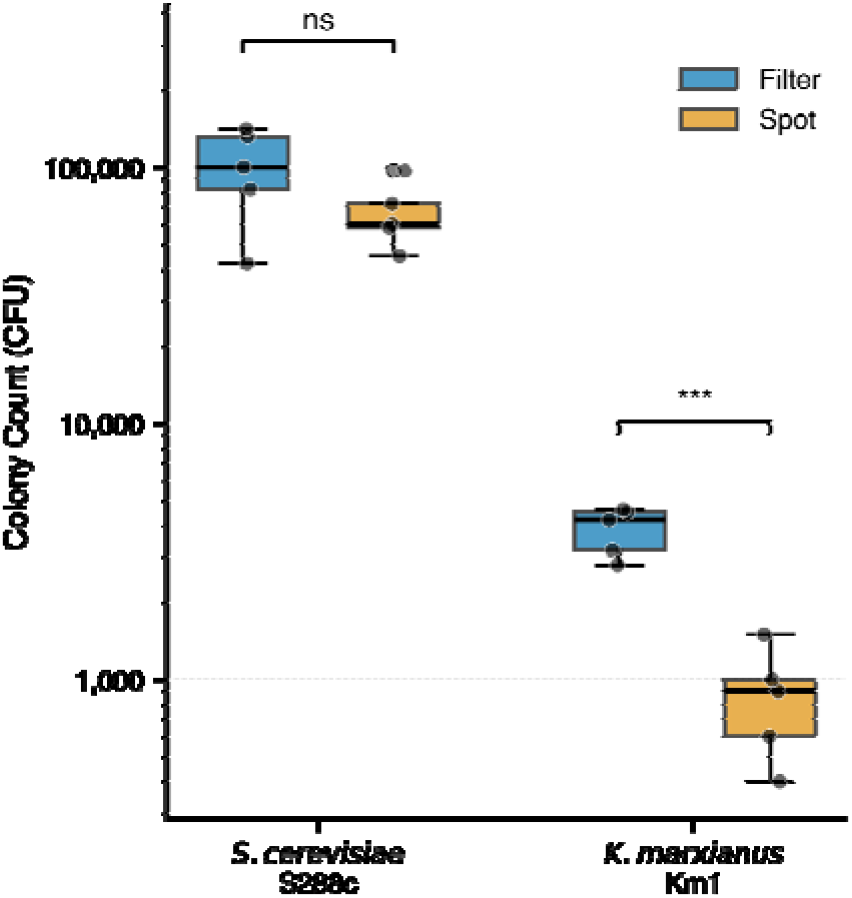
Comparison of filter and spot methods in *S. cerevisiae* and *K. marxianus*. Transformation results in terms of total number of CFUs from the filter and spot-plating methods. Results were from 1 mL of 1 OD culture for both *S. cerevisiae* and *K. marxianus*. *** p < 0.001 Welch’s t-test.

Another five phylogenetically diverse ascomycete yeasts were tested with the unfiltered spot-plating method: *Metschnikowia pulcherrima, Pichia pastoris, Ogataea parapolymorpha, Lipomyces tetrasporus*, and *Botryozyma nematodophila*. To our knowledge, only *L. tetrasporus* has been transformed via AMT. Members of the other genera have been transformed with alternative methods (i.e. PEG-mediated protoplasting, lithium acetate chemical transformation, or electroporation) (23–26). Notably, genetic transformation of *Botryozyma spp*. has not been previously reported (Figure 4A and 4B). Nourseothricin was found to be an effective antifungal against all species, so each was transformed with variations of the nourseothricin resistance (*nrsR*) gene with different promoters: the wildtype *Streptomyces nourseii nrsR* (n) two codon optimized *nrsR* variants (on and ln), the *A. gossypii* TEF1 promoter (AT), and the *L. starkeyii* TRPC (LR) promoter.

**Figure 4:**
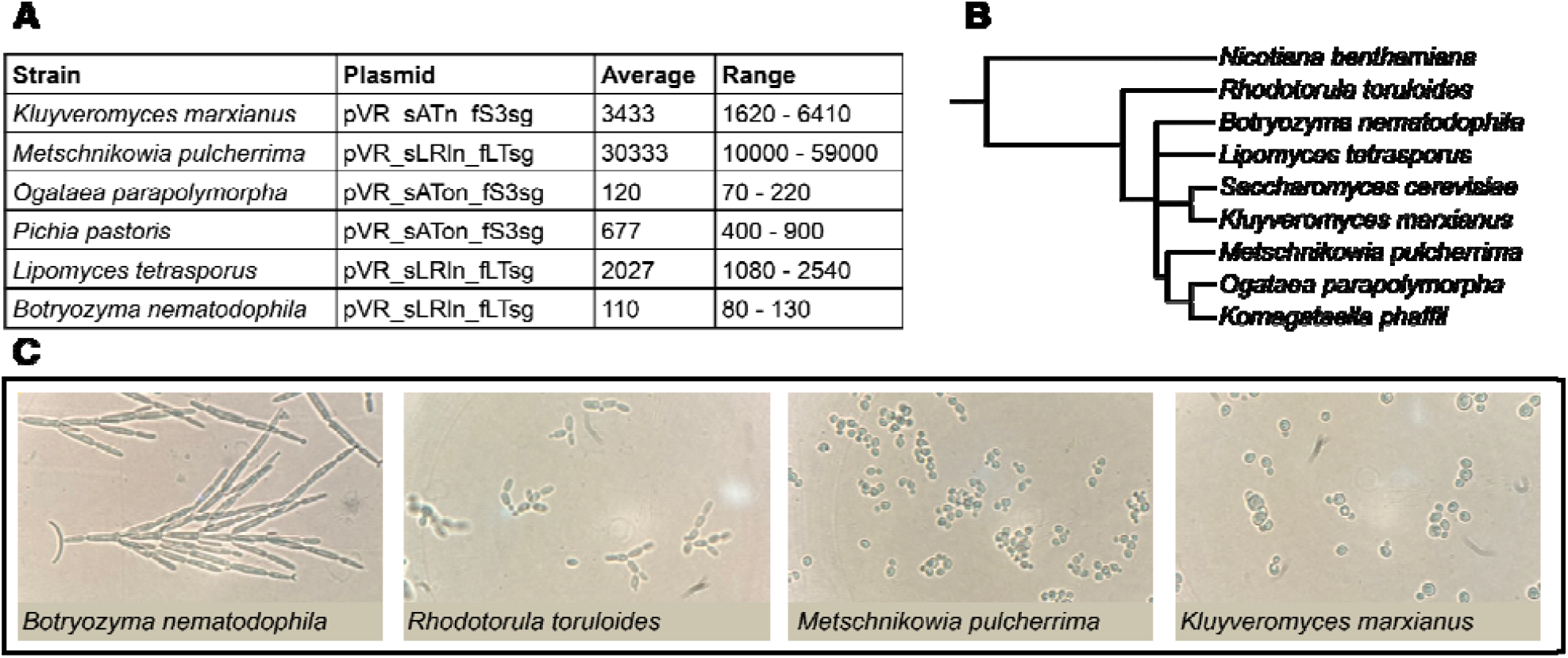
Transformation of diverse Ascomycete yeasts. **A.** Table of select transformation results for diverse Ascomycete yeasts. Complete table available in Table S3. All transformations were conducted using the spot-plating method with *Agrobacterium*:yeast OD·mL ratios of 1:1. **B**. Family tree of all tested microbes with *Nicotiana benthamiana* as an outgroup to root the tree. The tree was constructed via phyloT v2 using NCBI taxonomy (27). **C**. Microscopy illustrating the diverse morphologies of select tested microbes.

All variants of the selection cassette worked to varying degrees in all hosts except *M. pulcherrima* (Table S3). *M. pulcherrima* is a CTG-Ser yeast and could only be successfully transformed with the codon optimized *nrsR* cassettes, both of which lacked CTG codons (24). Transformation yields varied by species (from <100 to >100,000 colonies), possibly reflecting differences in morphology (Figure 4C), cell wall composition, T-DNA integration efficiency, selection marker expression, or preferred DNA repair mechanisms (HR vs NHEJ). Nonetheless, the fact that all seven species could be transformed via this protocol demonstrates that the unfiltered method is broadly applicable across diverse yeast lineages. While we simply quantified the total number of resulting transformants and not the efficiency, future work on host-specific optimization could evaluate this metric.

## Conclusions

The spot-plating method reported and optimized here streamlines AMT of non-model fungal species, a historically labor-intensive, time-consuming, and inconsistent process. By eliminating the vacuum and filter entirely, this protocol reduces hands-on time from several hours to approximately fifteen minutes for 12 transformations, notably increasing the speed and feasibility of AMT for large-scale functional genomic experiments. We furthermore found that this method improved transformation efficiency relative to prior filter plating protocols by up to 66% for *R. toruloides*. By systematic optimization of cell densities and resuspension volumes, we obtained more than 200,000 total transformants per transformation. This enables transformation with sufficient coverage for the construction of large libraries. Our successful transformation of seven yeast species spanning multiple genera using spot-plating confirms that the unfiltered approach is not limited to well-characterized model organisms but functions as a generalizable platform for onboarding molecular biology tools into non-model hosts. Successful transformations in *Botryozyma nematodophila* and *Metschnikowia pulcherrima* are of particular interest for their unique morphology and under-explored biotechnological potential (28–30). Despite its lower transformation efficiency relative to filter-plating in some hosts, spot plating is the more practical method, requiring far less hands-on time (15 minutes vs. 2 or more hours for 6 replicates) and enabling parallel processing by eliminating the need for vacuum pump apparatuses. While not iterated upon here, optimization of the unfiltered method can be easily conducted by researchers investigating new hosts, following the framework described here. This protocol brings large-scale functional genomics experiments in recalcitrant yeast species within practical reach for researchers who previously lacked the capacity to attempt them. The straightforward method of spotting a mixture of *Agrobacterium* and yeast onto an induction medium enables automated workflows, accelerating investigation and engineering in genetically recalcitrant yeasts.

## Methods

### Media, chemicals, and culture conditions

Bacteria were routinely propagated in Luria-Bertani (LB) medium (BD Biosciences, USA). Yeast were routinely propagated in yeast extract peptone dextrose (YPD) medium (Sigma-Aldrich, USA). *Escherichia coli* and *Agrobacterium tumefaciens* were grown at 37 °C and 30 °C, respectively. Liquid cultures were shaken at 200 RPM.

Antibiotics and concentrations used are as follows: kanamycin (50 mg/L, Teknova, USA), nourseothricin (50 or 100 mg/L, GoldBio, USA), G418 (300 mg/L, Sigma-Aldrich), carbenicillin (300 mg/L, Teknova, USA), cefotaxime (300 mg/L, Sigma-Aldrich). All other chemicals were purchased through Sigma-Aldrich.

Induction medium (IM) was composed as follows: 1 g/L NH_4_Cl, 300 mg/L MgSO_4_·7H_2_O, 150 mg/L KCl, 10 mg/L CaCl_2_, 0.75 mg/L FeSO_4_·7H_2_O, 144 mg/L K_2_HPO_4_, 48 mg/L NaH_2_PO_4_, 3.9 g/L 2-(N-morpholino)ethanesulfonic acid (MES) (RPI, USA), 2 g/L D-glucose, 400 µM acetosyringone, 10 mg/L thiamine hydrochloride, pH adjusted to 5.5 with KOH. 2% Difco Bacto Agar (BD Biosciences) was used for induction medium plates. Medium was autoclaved before the addition of filter-sterilized stock solutions of thiamine (100 g/L in H_2_O) and acetosyringone (400 mM in DMSO (Sigma-Aldrich)). We employ a high acetosyringone concentration (400 µM) which we previously found to be effective in *Rhodotorula toruloides* IFO 0880 (NBRC 0880). Researchers should adjust this for their yeast of interest as it likely affects transformation efficiency in different hosts (15,31).

### Strains and plasmids

All bacterial and yeast strains used in this work are listed in Table 1, and all plasmids are listed in Table 2. All plasmids generated in this paper were assembled using PCR and Gibson Assembly® and transformed into *E. coli* NEB-turbo using standard protocols (32). Primers were purchased from Integrated DNA Technologies (IDT, USA). Plasmids were isolated using the QIAprep Spin Miniprep kit (Qiagen, USA). Plasmid sequences were verified using whole plasmid sequencing (Plasmidsaurus, USA). *A. tumefaciens* was routinely transformed via electroporation as described previously (33).

**Table 1:** Strains employed in this study.

| Species | Source/Citation |
| --- | --- |
| *Agrobacterium tumefaciens* EHA105 | (34) |
| *Escherichia coli* NEBturbo | NEB |
| *Rhodotorula toruloides* NBRC 0880 | NBRC |
| *Saccharomyces cerevisiae* S288c | (35) |
| *Kluyveromyces marxianus* Km1 | (36) ABF_015686 |
| *Metschnikowia pulcherrima* 40-214 (ATCC 18406) | Phaff Collection, Davis, CA |
| *Botryozyma nematodophila* CBS 7442 | CBS, (28) |
| *Komagataella phaffia* (*Pichia pastoris*) X-33 | Invitrogen |
| *Ogataea parapolymorpha* DL-1 (ATCC 26012) | ATCC |
| *Lipomyces tetrasporus* NRRL Y-11562 (CBS 5910) | CBS, (37) |

**Table 2:**
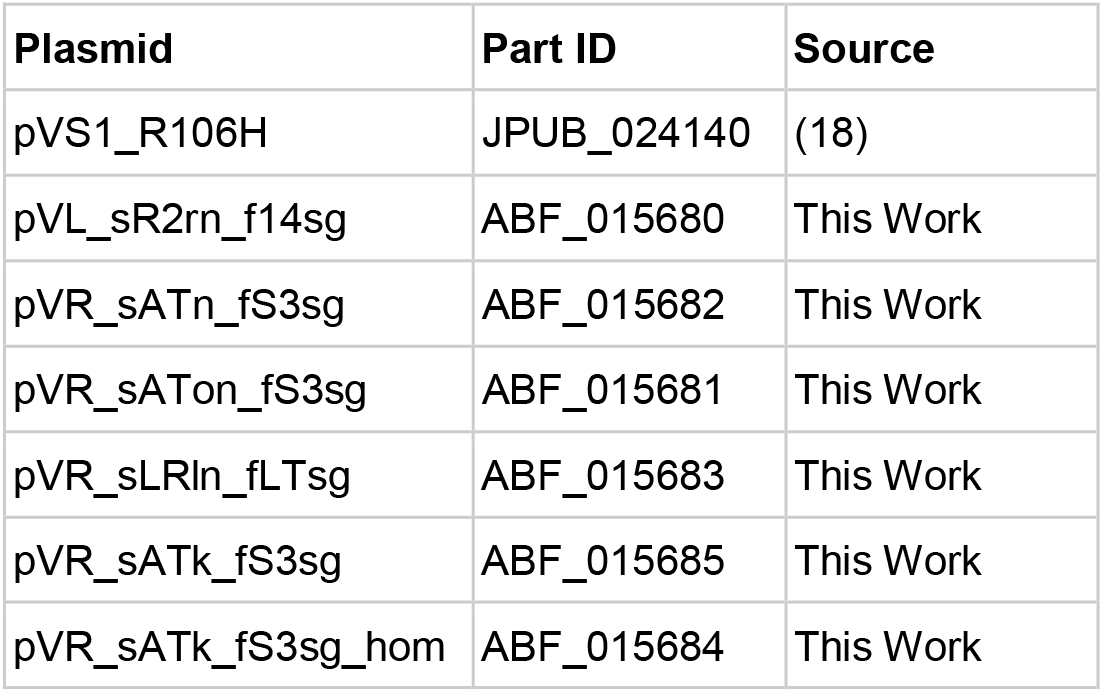
Plasmids used in this study. All available through public-registry.jbei.org and public-registry.agilebiofoundry.org.

### Vacuum filtration *Agrobacterium-*mediated transformation

This describes a protocol for up to 10 total transformations. This can be scaled up using larger culture volumes and centrifugation formats.

#### Day 1

A single colony of *Agrobacterium* carrying a binary vector was inoculated into a culture tube containing 10 mL LB medium with kanamycin and cultured overnight (∼16 hours, 30 °C, 200 RPM). The target yeast strain was inoculated from a single colony into a culture tube containing 10 mL YPD and grown in parallel.

#### Day 2

The overnight *Agrobacterium* culture was diluted to OD□□□ = 0.5 in 10 mL LB without antibiotics and grown at 30 °C with shaking until OD□□□ = 1.0 (∼2.5 hours). Cells were harvested by centrifugation at 5,000 × g for 15 minutes in 15-mL conical tubes (Falcon, Corning, USA), washed once with 10 mL induction medium (IM), and resuspended in fresh IM to a final OD□□□ = 1.0. The culture was then incubated at 30 °C with shaking overnight for *vir*-gene induction. Serial dilutions of the overnight yeast culture were prepared in fresh YPD so that one dilution would reach OD□□□ = 1.0 the following day.

#### Day 3

The OD□□□ of both the induced *Agrobacterium* and yeast cultures was measured. 1 mL of yeast culture (OD□□□ = 1.0) was transferred to a 1.5-mL microcentrifuge tube (Eppendorf, Germany) and pelleted at 5,000 × g for 5 minutes. The supernatant was aspirated, and the yeast pellet was resuspended in a volume of induced *Agrobacterium* culture yielding an equivalent OD□□□ ratio (e.g., 666 µL of OD 1.5 Agrobacterium culture). The combined cell suspension (1,000 µL) was applied in 250-µL increments onto a 0.45-µm mixed cellulose ester membrane filter (HAWP; MilliporeSigma, USA) under vacuum. The filter was transferred with sterile forceps, cell-side up, to an IM agar plate and incubated at 26 °C for 4 days.

#### Day 7

The filter was carefully removed from the IM plate with sterile forceps and placed into a 50-mL conical tube containing 5 mL of YPD. The tube was vortexed vigorously for 2 minutes to resuspend the cells. Serial dilutions were prepared, and aliquots were spread onto selective YPD agar plates supplemented with cefotaxime and carbenicillin to counter-select *Agrobacterium*. Plates were incubated at 30 °C for 2 days or until single colonies were visible.

### Unfiltered spot plating *Agrobacterium* mediated transformation

Days 1 and 2 were performed identically to the vacuum filtration protocol described above.

#### Day 3

The OD□□□ of both the induced *Agrobacterium* and yeast cultures was measured. 1 mL of yeast culture (OD□□□ = 1.0) was transferred to a 1.5-mL microcentrifuge tube (Eppendorf, Hamburg, Germany) and pelleted at 5,000 × g for 5 minutes. The supernatant was aspirated, and the yeast pellet was resuspended in the induced *Agrobacterium* culture at an equivalent OD□□□ ratio (i.e., 1:1 by cell density). The combined suspension was then centrifuged at 10,000 × *g* for 5 minutes, the supernatant was aspirated, and the cells were resuspended in 20 µL of sterile deionized water. The concentrated cell mixture was spotted directly onto the surface of an IM agar plate and incubated at 26 °C for 4 days.

#### Day 7

The cell mixture was scraped from the surface of the IM agar using a 10 µL sterile loop (Thermofisher Scientific, USA) and resuspended in 1 mL YPD by swirling. Cell suspensions were vortexed until cell clumps were no longer visible, serial dilutions were prepared, and aliquots were spread onto selective YPD agar plates supplemented with cefotaxime and carbenicillin to counter-select *Agrobacterium*. Plates were incubated at 30 °C for 2 days or until single colonies were visible.

### Microscopy and Spectrophotometry

For all cultures, OD□□□ was measured using a Spectramax M2 (Molecular Devices, USA). *Metschnikowia pulcherrima* and *Pichia pastoris* were prone to forming flocs in liquid culture, confounding optical density measurements. For accurate measurements, these cultures were diluted into an EDTA solution (50 mM, pH = 8) and vortexed to disrupt flocs before reading.

Colonies of yeast species were inoculated and cultured overnight in YPD (30 °C, 200 RPM). Cultures were spotted onto microscope slides (VWR) and examined under a VistaVision compound microscope (VWR, USA) with 10 x magnification eyepiece and 100 x magnification objective (1000 x total magnification). Images were taken with an iPhone 14 Plus (Apple, USA) main camera (12 MP, 26 mm, *f*/1.5 aperture) held to the eyepiece lens without software magnification.

## Acknowledgements

Sandia National Laboratories is a multi-mission laboratory managed and operated by National Technology and Engineering Solutions of Sandia LLC., a wholly-owned subsidiary of Honeywell International Inc., for the U.S. Department of Energy’s National Nuclear Security Administration under contract DE-NA0003525. The investigation of diverse fungi in this study was supported by breakthroughs obtained through the Laboratory Directed Research and Development Program of Lawrence Berkeley National Laboratory under U.S. Department of Energy Contract No. DE-AC02-05CH11231. This study is part of the Agile BioFoundry (https://agilebiofoundry.org) supported by the U. S. Department of Energy, Office of Critical Minerals and Energy Innovation, and was part of the Joint BioEnergy Institute (https://jbei.org), supported by the U.S. Department of Energy, Office of Science, Biological and Environmental Research Program through contract DE-AC02-05CH11231 between Lawrence Berkeley National Laboratory and the U.S. Department of Energy. This report was prepared as an account of work sponsored by an agency of the United States Government. Neither the United States Government nor any agency thereof, nor any of its employees, makes any warranty, express or implied, or assumes any legal liability or responsibility for the accuracy, completeness, or usefulness of any information, apparatus, product, or process disclosed, or represents that its use would not infringe privately owned rights. Reference herein to any specific commercial product, process, or service by trade name, trademark, manufacturer, or otherwise does not necessarily constitute or imply its endorsement, recommendation, or favoring by the United States Government or any agency thereof. The views and opinions of authors expressed herein do not necessarily state or reflect those of the United States Government or any agency thereof. The publisher, by accepting the article for publication, acknowledges that the U.S. Government retains a nonexclusive, paid-up, irrevocable, worldwide license to publish or reproduce the published form of this manuscript, or allow others to do so, for U.S. Government purposes. The Department of Energy will provide public access to these results of federally sponsored research in accordance with the DOE Public Access Plan (http://energy.gov/downloads/doe-public-access-plan). Funding for open access charge: U.S. Department of Energy. We thank Prof. John Dueber, Prof. Rachel Brem, and Dr. Ziyu Dai for generously providing diverse yeast strains and nourseothricin selection cassettes.

## Contributions

Conceptualization, P.A.A., M.R.I., M.G.T.; Methodology, P.A.A., M.R.I., M.G.T., M.J.S., A.L., G.M.G.; Investigation, M.R.I., M.G.T., M.J.S., G.M.G., A.L.; Writing – Original Draft, M.R.I., M.G.T., A.L.; Writing – Review and Editing, All authors; Resources and supervision, M.G.T., D.L., J.M.G., P.M.S.

## Competing Interests

M.G.T., M.J.S., and P.M.S. have financial interests in BasidioBio.

## Notes

### Summary of Updates

The manuscript, now titled “A membrane-free spot-plating protocol for Agrobacterium-mediated transformation of diverse yeasts,” underwent revision between submissions. The original title, which used the word “Unfiltered” and specified basidiomycete and ascomycete yeasts, was replaced to better reflect the method and broaden scope The abstract was substantially rewritten. A sentence explaining the Agrobacterium-mediated transformation mechanism was added. The framing shifted from presenting a novel method to contextualizing it within prior literature. The bibliography changed from a Nature-style format to Vancouver format for Scientific Reports. The reference count grew from 25 to 37, with twelve new references supporting added mechanistic context, historical precedents, and phylogenetic methods.

