## Supplementary figures and images for "A membrane-free spot-plating protocol for *Agrobacterium*-mediated transformation of diverse yeasts"

### Supplemental Figure 1

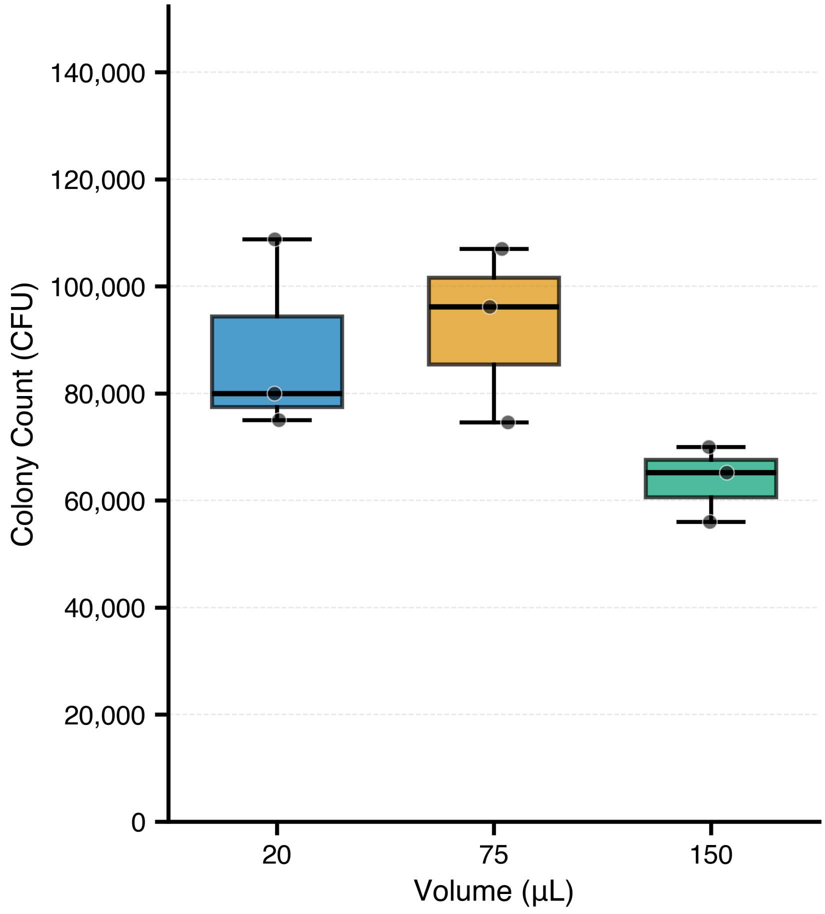
